# Chaperones Hsc70 and Hsp70 Play Distinct Roles in Bocaparvovirus Minute Virus of Canines Infection

**DOI:** 10.1101/2022.09.19.508470

**Authors:** Jianhui Guo, Jinhan Sun, Yan Yan, Kai Ji, Zhiping Hei, Liang Zeng, Huanzhou Xu, Xiang Ren, Yuning Sun

**Affiliations:** Department of Biochemistry and Molecular Biology, School of Basic Medical Science, Ningxia Medical University, Yinchuan, 750004, China; Department of Clinical Medicine, School of Clinical Medicine, Ningxia Medical University, Yinchuan, 750004, China; Center for Emerging Infectious Diseases, Wuhan Institute of Virology, Chinese Academy of Sciences, Wuhan, Hubei, 430071, China; University of Chinese Academy of Sciences, Beijing, 100049, China

**Keywords:** bocavirus, Hsc70, Hsp70, MVC, virus infection

## Abstract

Minute virus of canines (MVC) belongs to the genus *Bocaparvovirus* and reproduces rapidly in its permissive cells Walter Reed/3873D (WRD). The crosstalk between MVC and WRD is poorly characterized in terms of molecular requirements and mechanisms. Here, we identified two novel cellular proteins Hsc70 and Hsp70 that interact with both NS1 and VP2 via mass spectrometry (MS), co-immunoprecipitation (Co-IP) and confocal immunofluorescence assays (IFA). Hsp70 was upregulated upon MVC infection. Respective silencing of Hsc70 and Hsp70 led to contrasting results at nearly every stage of MVC life cycle, including virus entry, transcription, translation, replication and production. Strikingly, transfection with low and high dose of pFlag-Hsp70 contributed to opposing impacts on viral protein levels and virus production possibly through a ubiquitin-dependent manner, indicating that MVC is quite sensitive to the levels of Hsp70. Treatment with quercetin and VER155008, two Hsp70 family inhibitors, both significantly decreased viral replication and particle levels. Together, these results illustrated that both Hsc70 and Hsp70 are involved in MVC life cycle, and targeting to Hsp70 family may represent a novel anti-MVC mechanism.

## INTRODUCTION

Minute virus of canines (MVC), also known as canine bocavirus, belongs to the genus *Bocaparvovirus* which is newly established in the subfamily *Parvovirinae* within the *Parvoviridae* family (Binn, Lazar et al., 1970). Infection of MVC causes respiratory, cardiac and gastrointestinal symptoms in neonatal canines and infertility as well as stillbirths in pregnant bitches (Pollock & Coyne, 1993). Another member in *Bocaparvovirus* genus, human bocavirus 1 (HBoV1) which causes lower respiratory tract infections in children, exerts similar molecular characteristics to MVC. An infectious molecular clone of the HBoV genome (pIHBoV) has already been constructed and replicates in human embryonic kidney 293 cells (HEK293) (Shao, Shen et al., 2021). Recent years, HBoV1 was found able to infect well-differentiated/polarized primary or immortalized human airway epithelium (HAE) cultured at an air-liquid interface (HAE-ALI) (Deng, Yan et al., 2013), however, further research was limited due to the complexity of the ALI cultures. In our previous study, we have cloned and sequenced the full-length MVC genome, constructed the recombinant infectious clones of MVC (pI-MVC), and obtained invaluable information about DNA replication in its permissive cells, WRD (Walter Reed canine cell/3873D) cells (Sun, Chen et al., 2009).

MVC is a non-enveloped autonomous virus containing a very small single-stranded DNA genome. It is organized into multiple non-structural-encoding gene units in the left half and capsid-encoding gene units in the right. The whole genome contains three open reading frames (ORFs), of which the left ORF encodes the non-structural protein NS1 (~85/66kDa), the middle ORF encodes the non-structural protein NP1 (~20kDa), and the right ORF encodes three structural proteins: VP1 (~81kD), VP2 (~67/63kD), VP3 (~61KD). VP3 is the product of post-translational proteolytic cleavage during formation of VP2 and only exists in intact virions containing viral DNA (vDNA). VP2 is the main structural protein of viral capsid (90%) and is a key factor in determining host range (Li, Zhang et al., 2014). The large NS protein NS1 is a multifunctional protein essential for viral replication. Replication of MVC DNA of the NS1(-) mutant, of which the NS1 open reading form (ORF) was terminated early in pI-MVC, was totally abolished. Termination of a middle ORF NS protein NP1 in pI-MVC also reduced replication of MVC DNA by 320-fold (Sun et al., 2009). Studies using *Protoparvovirus* minute virus of mice (MVM) have also shown NS1 aided viral genome to establish and amplify its lytic infection in the MVM replication centers, which colocalized with cellular sites of DNA damage induced by viral infection. And NP1 was found to govern access to the capsid gene by promoting splicing and restraining internal polyadenylation of pre-mRNAs (Majumder, Wang et al., 2018). As parasitic organisms, efficient viral propagation is utterly dependent on host factors, including adsorption, disassembly, transcription, translation, replication and virus particle packaging (Flatt & Butcher, 2019). Host cells are stimulated to defend virus infection while forced to provide favorable environment for virus proliferation. Accordingly, host factors may either play positive or negative roles in viral life cycle.

The heat shock proteins (Hsps), which not only facilitate protein folding, assembly, transportation, but also assist misfolded proteins to refold into functional states and degrade impaired proteins, are crucial in maintaining cellular homeostasis (Hu, Yang et al., 2022). Based on these salient characteristics, Hsps, including Hsp70 family, are often usurped by viruses to benefit their own survival (Zhang & Yu, 2022). Hsp70 family is highly conserved upon sequence and structure. It is mainly composed of two functional domains. The nucleotide binding domain (NBD, 45kDa) at N-terminus with a V-shaped structure consists of two subdomains containing ATP binding sites, which possess ATPase activity, and the substrate binding domain (SBD, 25kDa) at the C-terminus. When proteins are misfolded, the exposed hydrophobic residues are recognized by SBD and degraded by ubiquitin-dependent systems (Kityk, Kopp et al., 2012, Rosenzweig, Nillegoda et al., 2019). Hsp70 isoforms (Hsp70s) are abundant within cells and can be represented by constitutively (Hsc70, 7lkDa) and highly inducible expressed (Hsp70, 70kDa) species, which share about 86% homology in human (Fernández-Fernández & Valpuesta, 2018, Rosenzweig et al., 2019) and 95% in canines. Hsp70 is slightly expressed under unstressed conditions but highly induced under stress conditions, such as virus infections and cancer diseases, while Hsc70 remains unchanged or is slightly upregulated (Fernández-Fernández & Valpuesta, 2018). The inducible Hsp70 is functionally supplementary to protect cells from damage caused by external stimuli. The proteins in this family are evolutionarily conserved but functionally diversified. Though with high similarity, there are still distinct differences between Hsc70 and Hsp70, for example, Hsc70 contains introns while Hsp70 does not, which ensures rapid and abundant expression of Hsp70 once transcription starts(Liu, Yang et al., 2014).

Interactions between Hsp70s and viral proteins have been shown to facilitate different stages of the viral life cycle. It has been widely reported that Hsp70s not only affect the replication of viruses, but also have important impacts on their transcription, protein expression and other processes. For example, Hsp70s promote protein synthesis and viral replication of adenovirus, assist in the transportation of proteins into the nucleus in polyomavirus and enhance viral replication of measles virus (Chromy, Pipas et al., 2003, Glotzer, Saltik et al., 2000, Wan, Song et al., 2020). NBD inhibitor JG40, an inhibitor of Hsp70s, significantly inhibits Zika virus (ZIKV) propagation when added during entry and post-entry steps, suggesting that Hsp70s are required not only at the initial stage of infection, but also at the stages of viral RNA synthesis and infectious virions production (Pujhari, Brustolin et al., 2019, Taguwa, Yeh et al., 2019). DENV exploits Hsp70s in virus entrance, replication and virion production (Reyes-Del Valle, Chávez-Salinas et al., 2005, Taguwa, Maringer et al., 2015).

In this study, we performed affinity purification of NS1/VP2-interacting proteins, identified them through mass spectrometry experiments and found the NS1/VP2-Hsp70s connections in infected cells. We here show that knockdown of Hsc70 contributed to significant reduction at nearly every stage of infection, including viral entry, protein synthesis, replication and production. While knockdown of Hsp70 generated promotive effect on virus proliferation. In addition, low and high dose of pFlag-Hsp70 transfection (2μg, 6μg) seemed to exert opposing impacts on viral protein levels and virus production. Moreover, we proved that Hsp70 regulates viral proteins through a ubiquitination-dependent pathway. Importantly, two inhibitors of Hsp70 family, quercetin and VER155008, significantly and efficiently suppressed virus replication and production, which supports the concept that Hsp70s inhibitors can provide effective therapeutic strategy for MVC infection. Thus, our results illustrated that Hsc70 and Hsp70 are directly required for MVC life cycle, and targeting for Hsp70s may represent a novel antiviral mechanism.

## RESULTS

### Both Hsc70 and Hsp70 interact with MVC NS1 and VP2, respectively

Virus infection involves the participation of various host factors. To screen the novel host factors that interact with viral proteins, a combination of immunoprecipitation and mass spectrometry (MS) analyses was performed. First, the MVC-infected WRD cells were lysed and immunoprecipitated with anti-NS1 and anti-VP2 antibodies. Proteins binding to NS1 and VP2 were pulled down and analyzed by SDS-PAGE followed by Coomassie blue staining. As shown in Fig.1A, NS1, VP2 and IgG samples were screened via MS analysis. Compared to the IgG control, 18 proteins were discovered to interact with both NS1 and VP2. Proteins in NS1 and VP2 groups were ranked according to their numbers of unique peptides (>4) and interestingly, both of Hsc70 (HSPA8) and Hsp70 were found in each group and involved in nucleotide binding via Gene Ontology (GO) analysis (https://david.ncifcrf.gov/list.jsp) (GO:0000166), because of which we chose for the subsequent research.

**Figure 1.**
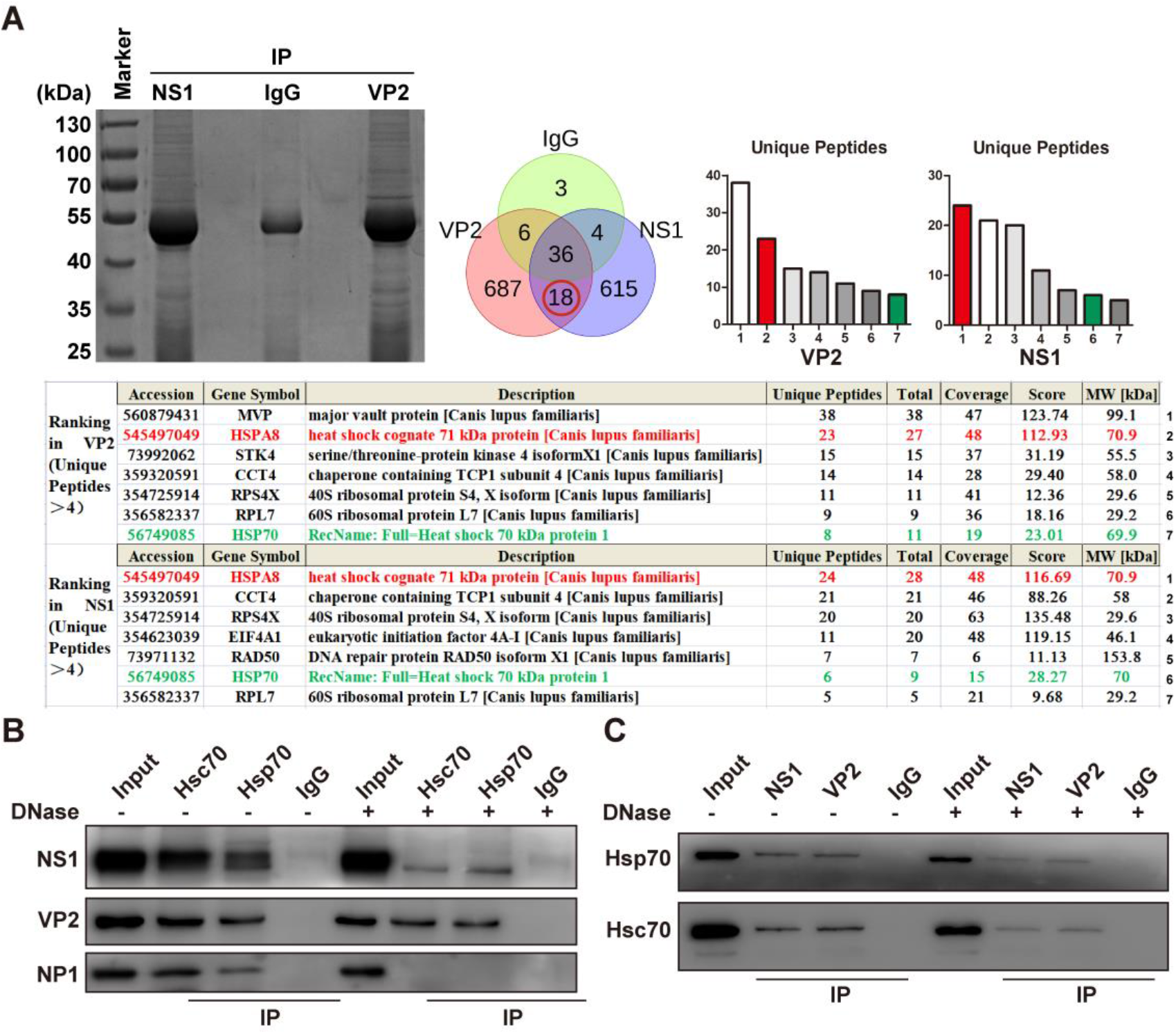
Screening and verification of host cells interacted with viral proteins. A WRD cells infected with MVC for 48 h were collected and lysed in NP40 buffer for 30 min on ice. Respective NS1 and VP2 antibodies as well as protein A/G beads were added to the cell lysis according to the manufacture’s instruction. IgG antibody was used as a negative control. Samples were subject to SDS-PAGE and Coomassie blue staining and then analyzed with mass spectrometry. Predicted cellular proteins that interacted with both NS1 and VP2 were screened by Bioladder based on IgG control data (https://www.bioladder.cn/web/#/pro/index). Rankings in NS1 and VP2 groups were based on unique peptides (>4). B Western blot analyses of Hsc70, Hsp70 and NS1, VP2, NP1 immunoprecipitations. Infected WRD cells were processed as lysed and co-immunoprecipitation (Co-IP) assays were performed with or without DNase as described. Hsc70 and Hsp70 antibodies and protein A/G beads were added to the cell lysis according to the manufacture’s instruction. IgG antibody was used as a negative control. Interactions between Hsc70, Hsp70 and NP1 vanished but still exist with NS1 and VP2. C Western blot analyses of NS1, VP2 and Hsc70, Hsp70 immunoprecipitations. Infected WRD cells were lysed and dealt with or without DNase. NS1 and VP2 antibodies and protein A/G beads were added to the cell lysis. Interactions between NS1, VP2 and Hsc70, Hsp70 could be detected.

To confirm the interaction between MVC and Hsp70s in cells, we performed Co-IP assays in WRD cells infected with MVC. Infected cells were lysed and co-immunoprecipitated using anti-Hsc70 and -Hsp70 antibodies for Western blotting using anti-NS1, -VP2 and -NP1 antibodies. With or without DNase, the whole cell lysates and the IgG sample were used for input and negative control, respectively. As shown in Fig. 1B, both MVC NS1 and VP2 formed complexes with cellular Hsc70 and Hsp70 in infected cells whether the DNase was present or not, while interactions between NP1 and Hsc70, Hsp70 vanished under the effect of DNase, indicating that the interaction between Hsc70/Hsp70 and NS1/VP2 was not mediated by vDNA. This was further validated by mutual Co-IP assays using anti-NS1 and -VP2 antibodies (Fig. 1C). Next, we verified these interactions in cos-1 cells co-transfected with plasmids expressing Hsc70 or Hsp70 and NS1 or VP2 followed by mutual Co-IP assays. As shown in Fig. 2A, protein-protein interactions were observed in cells co-transfected with pHA-NS1 and pFlag-Hsc70/-Hsp70. A parallel experiment was performed in which cos-1 cells were transiently co-transfected with pCDNA-VP2 and pFlag-Hsc70/-Hsp70 and similar results were obtained (Fig. 2D). To further characterize the interactions between Hsc70, Hsp70 and MVC NS1, VP2, we generated transiently expressed plasmids of Hsc70 and Hsp70, of which nucleotide binding domain on the N terminal (N) and substrate binding domain on the C terminal (C) were labeled with EGFP fusion respectively. Mutual Co-IP experiments demonstrated that both of the two domains of Hsc70 and Hsp70 interact with viral NS1 and VP2 (Fig, 2B, C; Fig. 2E, F). After confirming NS1/Hsc70 and NS1/Hsp70 interactions in vivo biochemically, we further investigated these interactions by performing immunofluorescence assay (IFA) in infected cells. WRD cells infected for 48 hours were then conducted by indirect IFA with antibodies against Hsc70/Hsp70 and NS1. As shown in Fig. 3, it was obvious that in uninfected WRD cells, Hsc70 localized in cytoplasm and nucleus (Fig, 3A), as well as Hsp70 but with less expression (Fig, 3B). Interestingly, they both translocated from cytoplasm into nucleus mostly after infection, and Hsp70 was markedly upregulated. Besides, as a viral replication-related protein, NS1 almost entirely aggregated in the nucleus as expected. Furthermore, their localizations were verified in cos-1 cells. pEGFP-Hsc70/-Hsp70 and pHA-NS1 were co-transfected into cos-1 cells, respectively. At 48 h post-transfection, cells were harvested with NS1 antibodies and DAPI and observed under confocal microscope. Consistent with above, Hsc70/Hsp70 were concentrated with viral NS1 (Fig. 3 A, B). Parallel experiments were carried out in which cos-1 cells co-transfected with pCDNA-VP2 and pEGFP-Hsc70/-Hsp70. Differently, VP2 almost entirely colocalized with Hsc70 and Hsp70 in cytoplasm (Fig. 3C, D). These results are sturdy evidence that Hsc70/Hsp70 coopt processing with NS1 and VP2 in cells.

**Figure 2.**
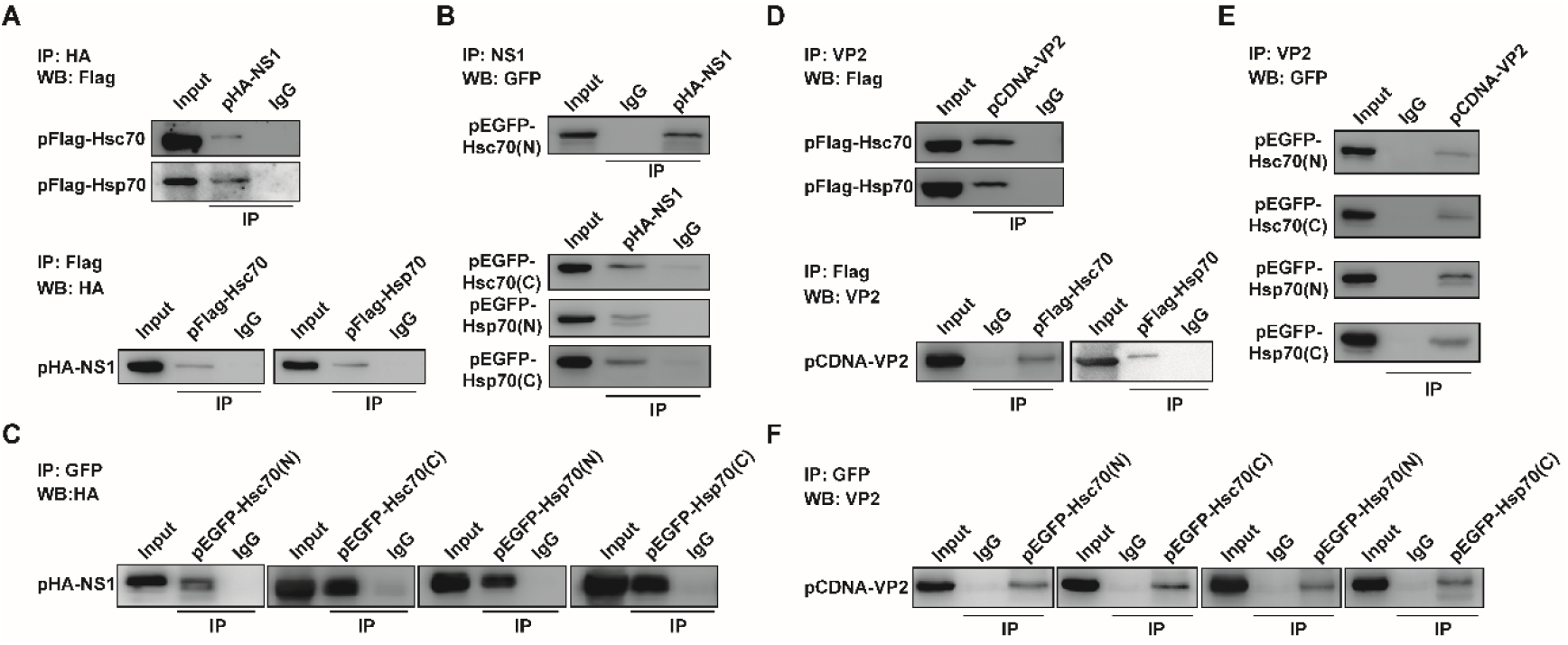
Hsc70, Hsp70 bind NS1, VP2 through both nucleotide binding domain (N) and substrate binding domain (C) A Plasmids expressing Hsc70/Hsp70 and NS1 were co-transfected into cos-1 cells respectively. Mutual co-IP experiments were performed and interactions between Hsc70/Hsp70 and NS1 were detected by Western blot. B, C Plasmids expressing Hsc70(N) / Hsc70(C) / Hsp70(N) / Hsp70(C) and NS1 were co-transfected into cos-1 cells respectively. Mutual co-IP experiments were performed and interactions between Hsc70/Hsp70 and NS1 were detected by Western blot. D Plasmids expressing Hsc70/Hsp70 and VP2 were co-transfected into cos-1 cells respectively. Mutual co-IP experiments were performed and interactions between Hsc70/Hsp70 and VP2 were detected by Western blot. E, F Plasmids expressing Hsc70(N) / Hsc70(C) / Hsp70(N) / Hsp70(C) and VP2 were co-transfected into cos-1 cells respectively. Mutual co-IP experiments were performed and interactions between Hsc70/Hsp70 and VP2 were detected by Western blot.

**Figure 3.**
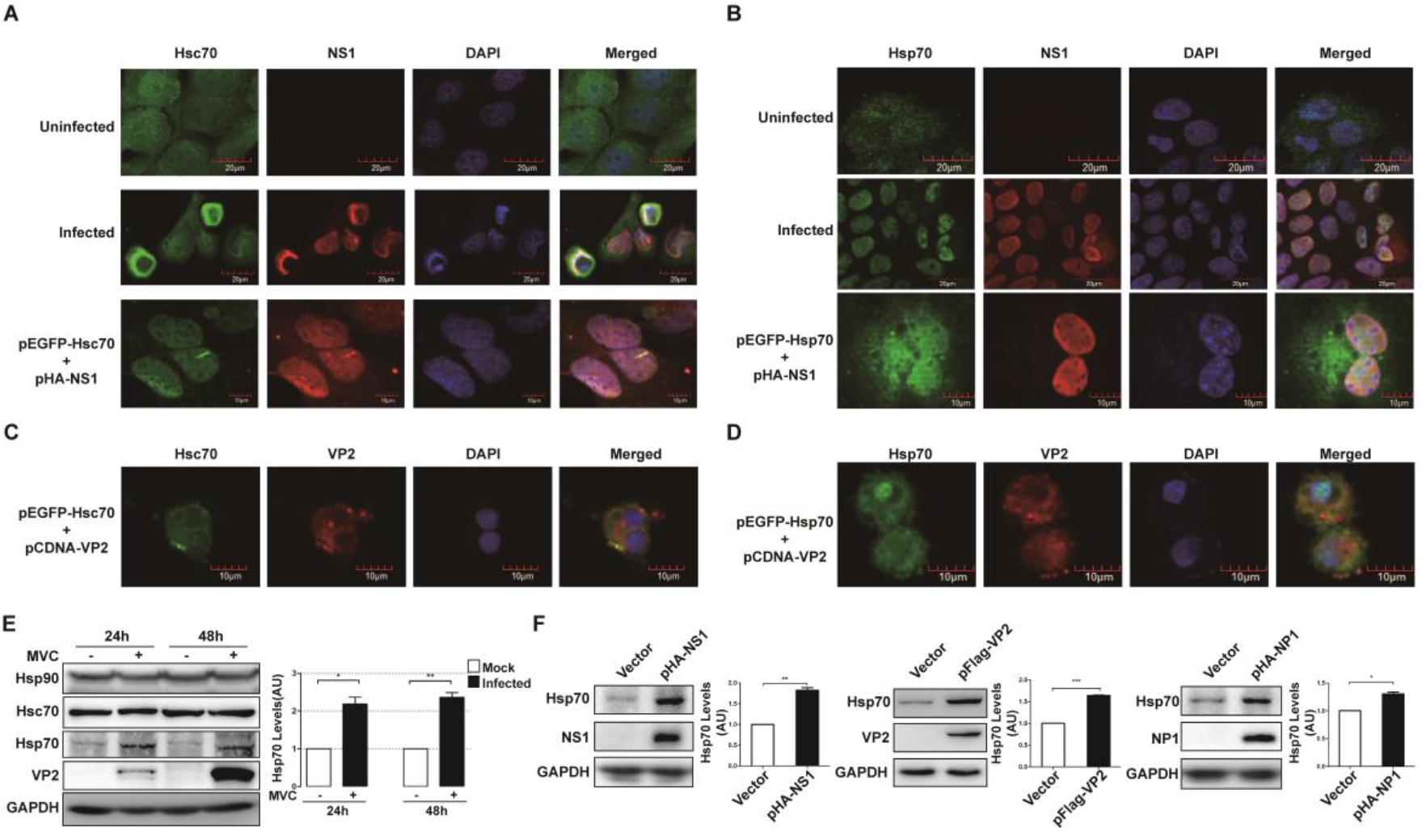
Localization of Hsc70, Hsp70 and NS1, VP2 and Hsp70 expression was elevated after MVC infection. A, B IFA analyses of the localization of Hsc70, Hsp70 and NS1. Top panels: IFA of uninfected WRD cells were used as negative control; middle panels: localization of Hsc70, Hsp70 and NS1 in infected WRD cells; bottom panels: localization of transcriptional Hsc70, Hsp70 and NS1 in cos-1 cells. C, D Localization of transcriptional Hsc70, Hsp70 and VP2 in cos-1 cells. Transient expression of EGFP fusion was used to label Hsc70 (pEGFP-Hsc70) and Hsp70 (pEGFP-Hsp70) as described. E Left panels: proteins in infected WRD cells were extracted at 24 hpi and 48 hpi and Hsc70, Hsp70 and Hsp90 accumulations during MVC infection were analyzed by Western blot. Right panels: Quantitative analyses of Hsp70 accumulations normalized to GAPDH levels (n = 3 independent experiments). F Top panels: Western blot analysis of Hsp70 levels after transfections of viral protein-expressing plasmids. Bottom panels: densitometry of the Western blot in top panels. Hsp70 levels were calculated using Image J software (n = 3 independent experiments). Data information: In (E, F), error bars represent the standard error of the means (SEM). Asterisks indicate statistically significant differences between the indicated groups (paired t-test, \**P*<0.05, \*\**P* < 0.01, \*\*\**P* < 0.001).

### Hsp70 is upregulated after MVC infection

IFA results above have clearly shown an obvious increase in Hsp70 at 48 h post-infection (hpi), which might be related to its stress-inducible characteristic under stress conditions, such as virus infection. We further confirm this upregulation in MVC-infected cells at different time points post-infection by Western blot (Fig. 3E). An increased Hsp70 level was observed in MVC-infected cells, as compared to the mock group; whereas Hsc70 nearly remained unchanged as expected. It should be noted that this Hsp response was specific, since the levels of Hsp90 were not increased upon viral infection. Levels of Hsp70 were also tested in WRD cells transfected with virus protein-expressing plasmids, pHA-NS1, pFlag-VP2 and pHA-NP1, respectively (Fig. 3F). Hsp70 levels were seen significantly upregulated under the circumstances of transfection of each plasmid. These data clearly indicated that the expression of intracellular Hsp70 was stimulated upon MVC infection, which might be in connection with viral proteins expressions. Considering the interactions between NS1, VP2 and Hsc70, Hsp70, and the homology between Hsc70 and Hsp70, we asked whether Hsc70 and Hsp70 participated in MVC life cycle. Thus, we next explored the respective roles of Hsc70 and Hsp70 in MVC life cycle.

### Knockdown of Hsc70 suppresses MVC infection in persistently infected WRD cells

We utilized short hairpin RNA (shRNA) against Hsc70 to determine its function in MVC infection. Knockdown efficiency of shHsc70 was examined, interestingly, an increasement in Hsp70 expression was observed in Western blot analysis when silencing Hsc70, with a higher increase in Hsp70 expression under the condition of relatively less Hsc70 (Fig. 4B). We speculated that Hsp70 was manipulated by the cellular compensatory mechanisms for maintaining proteostasis in the absence of Hsc70. The shHsc70-1 and/or shHsc70-2 were chosen for the subsequent experiments. As shown in Fig. 4C, qRT-PCR results of the relevant mRNAs established the efficiency of target Hsc70 knockdown. WRD cells treated with shHsc70s as well as their corresponding negative control shRNA (shNC) were infected with MVC, and viral proteins, replication, production and DNA levels determined. Knockdown of Hsc70 resulted in a significant decrease in virus protein levels (Fig. 4D) as detected by Western blot at 48 hpi. Surprisingly, shHsc70-2, which beard a lower knockdown efficiency, exerted a better inhibitory effect on MVC infection. In contrast, shHsc70-1 caused smaller though significant reduction, compared to shHsc70-2. As both viral non-structural and structural proteins, NS1 and VP2 were suppressed, similar results were observed accordingly, as performed by Southern blotting, qRT-PCR and focus-forming assay (FFA), in virus replication (Fig. 4E, F) and viral particle production (Fig. 4G). We speculated that the discrepancy of compensatory induced Hsp70 and/or other cellular factors due to deletion of a main molecular chaperone might mask a potential role for Hsc70 in viral infection. Therefore, we conclude that Hsc70 is directly involved in MVC propagation and the function of Hsp70 required further exploration.

**Figure 4.**
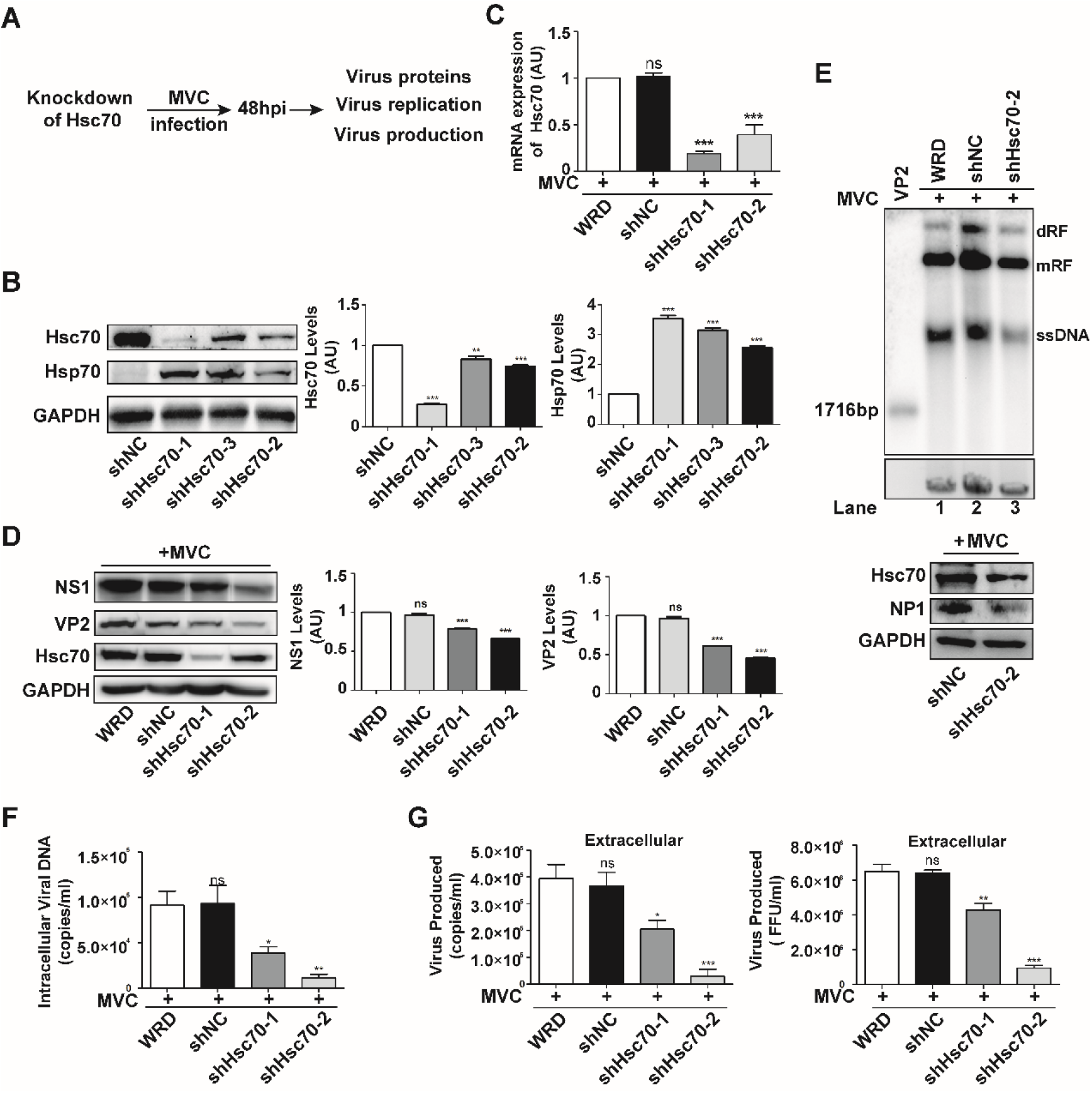
shRNA-mediated knockdown of Hsc70 significantly reduced MVC propagation. A Schematic diagram of Hsc70 knockdown experimental process. B, C Detection of silencing efficiency of Hsc70 by Western blot (B) and qRT-PCR (C) analyses. Hsp70 levels were examined simultaneously (B). D WRD cells were infected with short hairpin (sh) RNA-mediated (shHsc70-1, shHsc70-2) lentivirus and screened with puromycin (puro) at 1 μg/ml for at least 48 h. After MVC infection, NS1 and VP2 protein expressions were analyzed via Western blot. E Hirt DNA was extracted after virus infection as described. Production of replicative intermediates (dRF, dimmer replicative form; mRF, monomeric replicative form) and single-stranded DNA genome (ssDNA) was measured by Southern blot to assess viral replication in MVC-infected WRD cells. F qRT-PCR was performed to confirm the intracellular viral DNA loads to assess viral replication. G qRT-PCR and focus forming assay (FFA) experiments were performed to confirm the secreted virion loads. Data information: In (B-G), data shown in each panel are representative of three independent experiments. Statistical significance was determined with one-way ANOVA. Asterisks indicate significant differences (\**P* < 0.05, \*\**P* < 0.01, \*\*\**P* < 0.001). Error bars represent standard error of the means (SEM).

### Divergent effects of variant Hsp70 levels on viral propagation in infected cells

To assess the role of cytosolic inducible Hsp70 in the MVC life cycle, knockdown of Hsp70 experiments were performed then. Western blot and qRT-PCR analyses showed efficient elimination of Hsp70 in WRD cells (Fig. 5B, C), with shHsp70-4 exerting lower Hsp70 levels. Following 48 h infection of Hsp70 knockdown cells with MVC, we examined viral protein and mRNA expression levels simultaneously. Beyond our expectation, compared to normal and shNC groups, treatment with Hsp70 shRNAs did not decrease intracellular viral protein and mRNA levels; instead, statistically evident multiplications were observed in both shHsp70-4 and shHsp70-5 cells, with higher levels of infection in shHsp70-4 cells (Fig. 5B, C). In addition, qRT-PCR and FFA results suggested that vDNA replication and virus particle production were markedly heightened at 48 hpi which showed consistency with above results (Fig. 5D, E).

**Figure 5.**
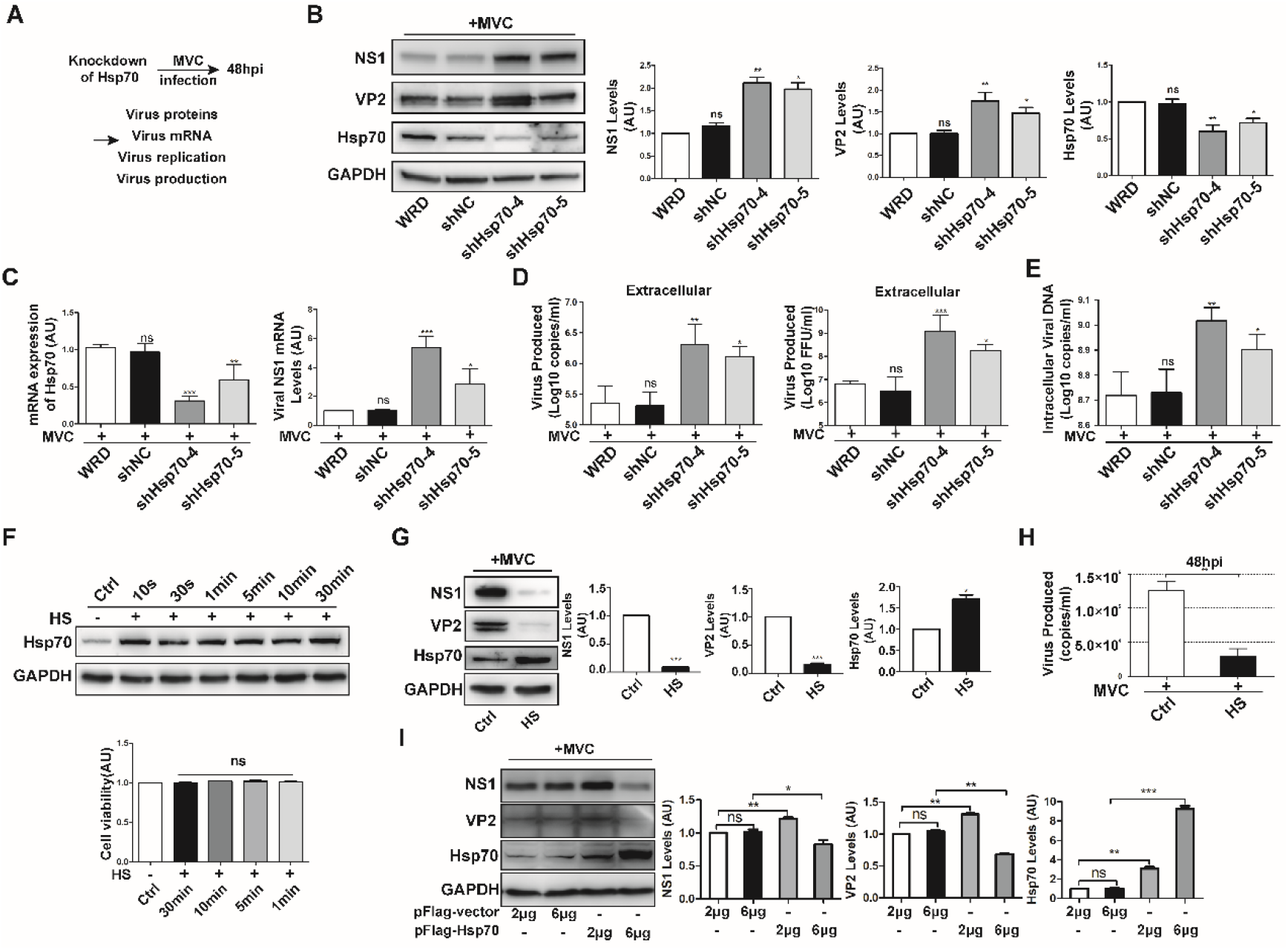
Hsp70 is essential for infection of MVC in WRD cells. A Schematic diagram of Hsp70 knockdown experimental process. B, C Detection of silencing efficiency of Hsp70 by Western blot (B) and qRT-PCR (C) analyses. WRD cells were infected with shRNA-mediated (shHsp70-4, shHsp70-5) lentivirus and screened with puro at 1 μg/ml for at least 48 h. After MVC infection, NS1 and VP2 protein expressions were analyzed via Western blot (B). Total RNA was extracted by trizol after virus infection and reverse transcribed to cDNA as described. qRT-PCR was performed to confirm viral mRNA expression (C). D qRT-PCR and focus forming assay (FFA) experiments were performed to confirm the secreted virion loads. E Intracellular viral DNA was extracted using a Quick-DNA^™^ Viral Kit (ZYMO Research) according to the manufacturer’s instructions and viral replication was assessed via qRT-PCR. F Top panels: Western blot analysis was performed to determine Hsp70 levels after HS treatment; bottom panels: WRD cells applied to heat shock (HS) treatment at 42 °C for different time were assessed by CCK-8 assays. G, H WRD cells were subjected to MVC infection after HS. NS1 protein expression and virus particles were extracted and detected at 48 hpi via Western blot and qRT-PCR. I WRD cells were transfected with different amounts of pFlag-Hsp70 plasmids (2μg, 6μg) and then infected with MVC. Western blot was performed to confirm levels of viral proteins. Data information: In (B-I), data shown in each panel are representative of three independent experiments. Statistical significance was determined with one-way ANOVA (B-F), paired t-test (G, I) and unpaired t-test (H). Asterisks indicate significant differences (\**P* < 0.05, \*\**P* < 0.01, \*\*\**P* < 0.001). Error bars represent standard error of the means (SEM).

To better understand the impact of Hsp70 in MVC infection, overexpression or heat shock (HS) treatment of Hsp70 was carried out in infected WRD cells by measuring MVC virus proteins and particle levels. HS was carried out at 42 °C and cell viability was measured upon four different HS durations. Results showed no detectable damage in WRD cells with HS treatment (Fig. 5F). When Hsp70 was overexpressed after HS, levels of MVC proteins (Fig. 5G) and virions (Fig. 5H) were all significantly decreased, compared with levels observed in the control groups. Next, we tested if overexpression via plasmids transfection has the same effect. The most striking and surprising results observed in infected WRD cells were the opposing levels of MVC proteins due to low and high dose of Hsp70 plasmids (2μg, 6μg) transfection. WRD cells were transfected with pFlag-Hsp70 24 h and subsequently infected with MVC for another 24 h. We see increased viral protein levels in the 2 μg group but decreased levels in the 6 μg group, as shown in Western blot analysis and blot quantification (Fig. 5I). These data indicated that MVC is extremely sensitive to the changes of intracellular Hsp70.

### Hsc70 and Hsp70 are involved in virus entry

Based upon the facts that some viruses utilize Hsp70 family proteins as cell receptors, and we have demonstrated the interactions between Hsc70, Hsp70 and MVC structural protein VP2, it is possible that Hsc70 and Hsp70 involve in the entry step of MVC infection. We first inspected the existence of Hsp70 and Hsc70 on WRD cell membrane. Expectedly, both of Hsc70 and Hsp70 were detectable on cytomembrane. Besides, we saw an adequate stimulation of cellular Hsp70 at 9 h post HS (Fig. 6A), a timepoint we chose for the followed MVC infection after HS.

**Figure 6.**
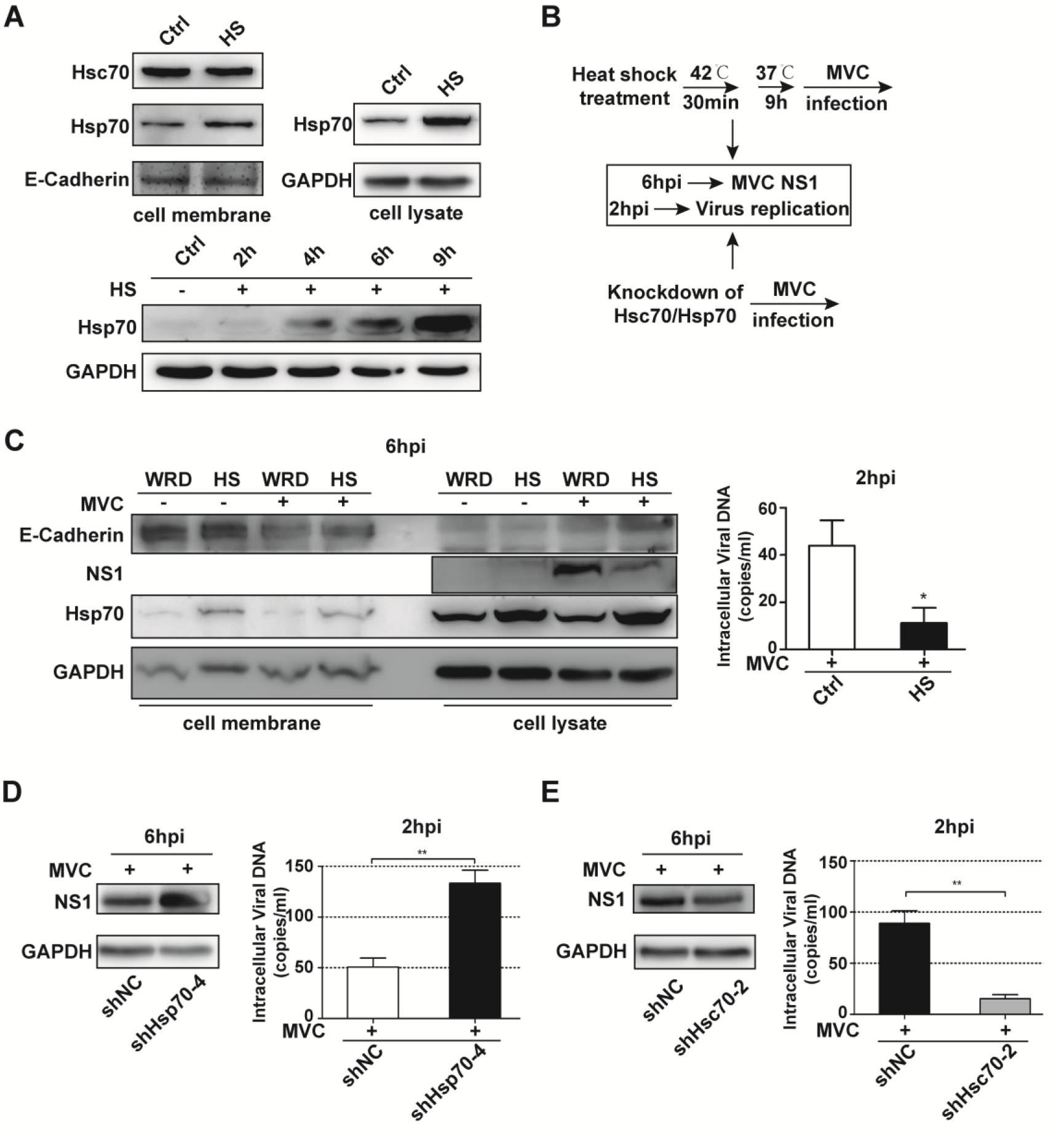
Hsc70 and Hsp70 participate in virus entry. A Top panels: WRD cells were treated with HS for 30 min and existence of Hsp70 and Hsc70 on cell membrane were detected. As expected, Hsp70 was upregulated both on cell membrane and in cytoplasm. Bottom panels: WRD cells were treated with HS for 30 min and then recover at 37°C. Western blot analysis was performed to confirm the time Hsp70 levels were dramatically induced during recovery, which would be used for MVC infection in this section. B Schematic diagram of experimental processes upon HS treatment and manipulation of Hsc70 and Hsp70. C WRD cells were treated with HS and recovered at normal temperature for 9 h followed by MVC infection. Western blot analysis was performed to confirm the virus entry loads at 6 hpi. qRT-PCR analysis was performed to confirm the amount of virus entry at 2 hpi. D, E qRT-PCR and Western blot analyses were performed to evaluate virus entry loads when individual silencing of Hsp70 (D) and Hsc70 (E). Data information: In (A, C-E), data shown in each panel are representative of three independent experiments. Statistical significance was determined with unpaired t-test (C-E). Asterisks indicate significant differences (\**P* < 0.05, \*\**P* < 0.01). Error bars represent standard error of the means (SEM).

WRD cells were subjected to MVC infection after HS treatment for 30 min and 37 °C culture for 9 h, at which point massive amounts of Hsp70 already existed. Viral NS1 levels at 6 hpi and vDNA at 2 hpi were extracted and measured to determine virus entry loads, as shown in Fig. 6C, prominent induction on cytomembrane Hsp70 has significantly suppressed virus entry. Hence, we sought to make certain that whether knockdown of Hsc70 and Hsp70 also affect the entry stage in MVC life cycle. shHsc70-2 and shHsp70-4 cells were subjected to virus infection followed by vDNA and proteins extraction at 2 hpi and 6 hpi, respectively. Knockdown of Hsc70 caused remarkably reduction while knockdown of Hsp70 contributed to enhancement in both extraction aspects (Fig. 6D, E). Conclusively, these observations strongly suggested that Hsc70 and Hsp70 are involved in MVC entry.

### Hsc70 and Hsp70 are also required for post-entry steps in MVC life cycle

Since that Hsc70 and Hsp70 participate in virus entry, we suspect the previous results of their intracellular function were due in part to their existence on cell membrane. In order to eliminate the interference of this aspect, WRD cells were infected with MVC prior to pFlag-Hsp70 transfection or HS treatment. Still, viral protein levels and virus production elevated in 2 μg group but reduced in 6 μg and HS groups (Fig. 7A, C). Besides, pI-MVC were transfected into shHsc70 and shHsp70 cells to bypass viral entry. NS1 levels decreased in shHsc70-2 cells and increased in shHsp70-4 cells compared with shNC both at 24 hpi and 48 hpi (Fig. 7B). These observations suggest that Hsc70 and Hsp70 have distinct roles in the viral life cycle both on viral entry and post-entry stages, i.e. viral protein translation, virus replication, and virion secretion.

**Figure 7.**
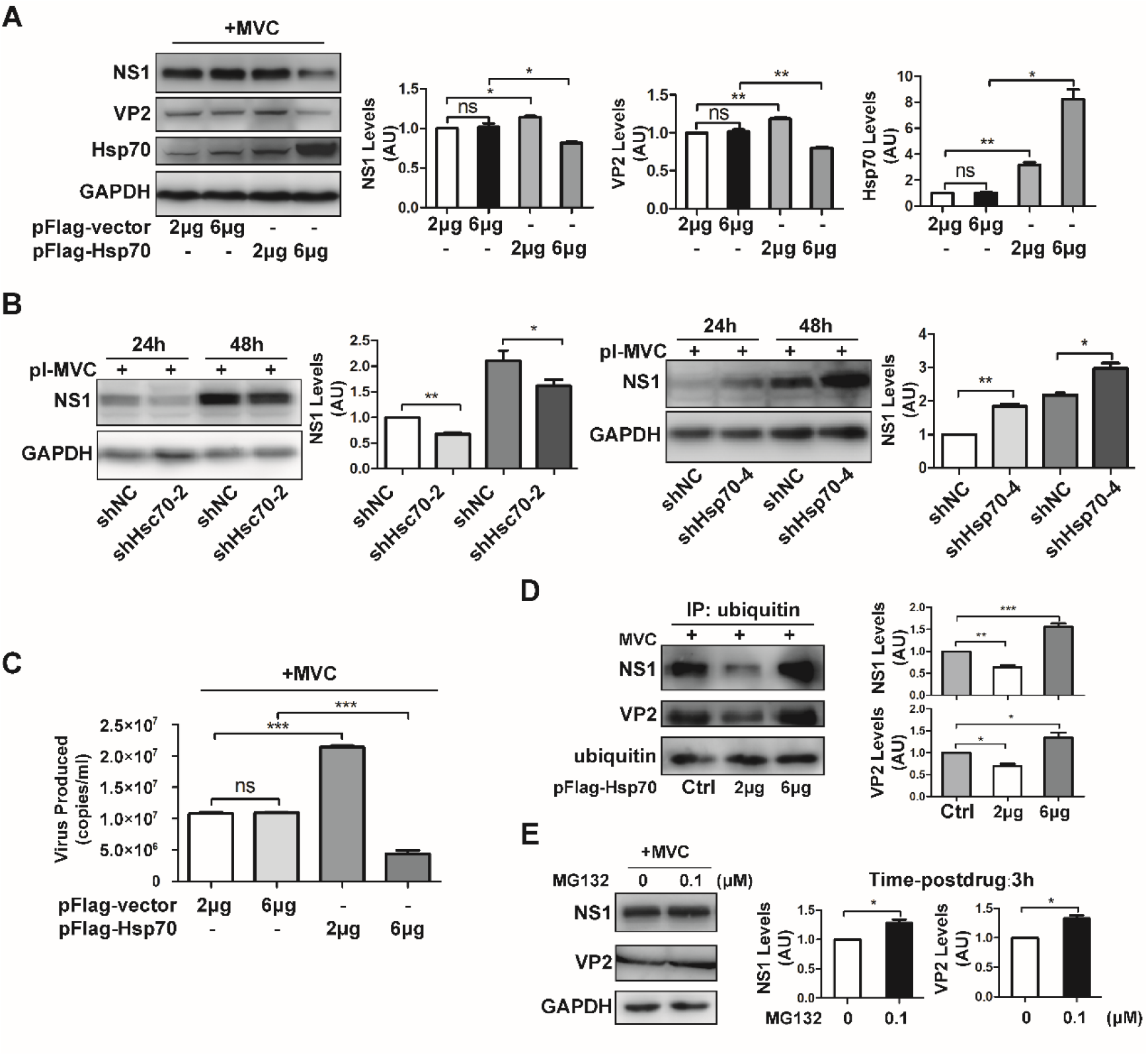
Hsp70 modulate viral proteins levels in post-entry steps through a ubiquitin-dependent degradation pathway in MVC life cycle. A, C WRD cells were infected with MVC and then transfected with different amounts of pFlag-Hsp70 plasmids (2μg, 6μg) or subjected to HS treatment. Western blot was performed to confirm viral proteins levels. qRT-PCR analysis was performed to determine virus production. B Bypass viral entry by pIMVC transfection. pIMVC was transfected into shHsc70 and shHsp70 cells respectively. D WRD cells were infected with MVC and then transfected with different amounts of pFlag-Hsp70 plasmids (2 μg, 6 μg). Infected lysates were subjected to IP using anti-ubiquitin antibodies. Immunopurified ubiquitinated proteins were subjected to Western blot analysis using anti-NS1, -VP2, and -ubiquitin antibodies as described. E WRD cells infected with MVC for 24 h were subjected to proteasome inhibitor treatment with MG132 (0.1 μM) or vehicle (0μM) for the last 3 h of infection. Cell lysates were analyzed by Western blot using anti-NS1 and -VP2 antibodies. Data information: In (A-E), data shown in each panel are representative of three independent experiments. Statistical significance was determined with unpaired t-test (C) and paired t-test (A, B, D, E). Asterisks indicate significant differences (\**P* < 0.05, \*\**P* < 0.01, \*\*\**P* < 0.001). Error bars represent standard error of the means (SEM).

### The amounts of intracellular viral proteins are associated with their ubiquitination

It has been widely reported that Hsp70 is a crucial regulator of CHIP-mediated ubiquitination and degradation (Wu, Wang et al., 2021). It is involved in the transfer of misfolded proteins to the proteasome system through a ubiquitination-dependent procedure. Thus, we speculated that the variation of viral protein levels connected with different amounts of Hsp70 is related to chaperone-dependent ubiquitination. To check our hypothesis, experiments were performed in which MVC-infected WRD cells were transfected with pFlag-Hsp70 (2μg, 6μg) and the ubiquitination levels of viral proteins were measured. Cell lysates were subjected to IP with anti-ubiquitin antibodies and viral proteins were detected via Western blot using anti-NS1/-VP2 antibodies. As shown in Fig. 7D, overexpression of 2μg Hsp70 was associated with a decrease of ubiquitinated viral proteins in infected WRD cells at 24 hpi, conversely, overexpression of 6μg Hsp70 was associated with an increase of ubiquitinated viral proteins. This is in agreement with the idea that viral proteins are transported by Hsp70 to a ubiquitinated degradation procedure.

To confirm our speculation, we employed a proteasome inhibitor, MG132, to see its impact on viral protein levels in infected WRD cells. WRD cells were infected for 24 h and then treated with MG132 for 3 hours. As illustrated in Fig. 7E, slight recover of viral protein levels was observed upon the addition of MG132 (0.1 μM), compared to the dimethyl sulfoxide (DMSO) vehicle (0 μM). Western blot analysis and blot quantification proved that viral proteins are normal substrates in the proteasome pathway.

### MVC infection is significantly impaired by inhibitors of Hsp70 family

To overcome the confounding effects of functional redundancy between Hsc70 and Hsp70, we employed two inhibitors of Hsp70s, VER155008 and quercetin, to further explore the function of Hsp70 family during MVC infection. The folding function of Hsp70s is coupled to its ATPase activity, hence, we took advantage of the allosteric inhibitor against this Hsp70-ATP interaction, VER155008. It competes with cellular ATP via inserting into the conserved ATP-binding pocket of Hsp70 family and therefore impairs the Hsp70 chaperone cycle. Another inhibitor quercetin is able to strongly suppress the synthesis of Hsp70s, with a more inhibitory tendency to the inducible Hsp70 (Ogawa, Sugawara et al., 2014). An advantage of these compounds over knockdown approaches is that they rapidly block the action of multiple Hsp70 isoforms. The activity of Hsp70 inhibitors on MVC replication was performed by time-of-addition assay and viral proteins expression were measured afterwards. Drugs-addition were sorted into four groups on the basis of adding time, as shown in Fig. 13A, of which both of the fourth (IV) groups exerted better inhibitory performance than the other three. Viral proteins were markedly repressed at 24 hpi (Fig. 8B, C). Hence, the follow-up drugs-addiction experiments were carried out using method IV (unless noted). In the presence or absence of drugs, the antiviral effect of these compounds was assessed by measuring their effects on viral mRNA levels monitored for up to 36 h. As expected, both inhibitors suppressed viral mRNA production to a similar extent in WRD cells (Fig. 8D). Drug interference of the compounds also drastically blocked cellular MVC replication (Fig. 8E) at different time points after infection, which indicates that successful establishment of intracellular viral replication state requires Hsp70s function. Extracellular particle production was dramatically reduced at 24 hpi to 48 hpi by qRT-PCR assays (Fig. 8F), especially in quercetin treatment.

**Figure 8.**
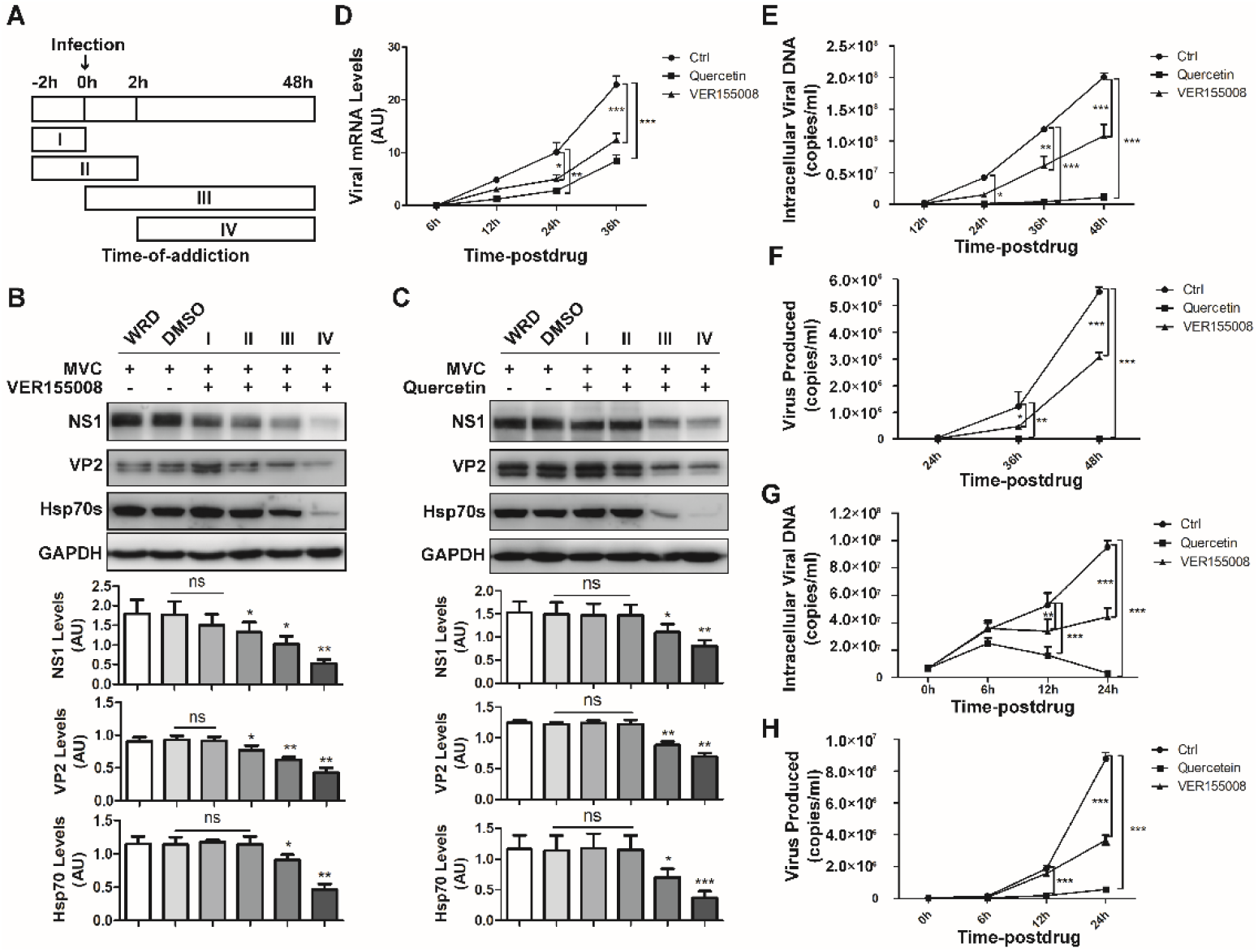
Quercetin and VER155008 markedly decreased MVC infection in many aspects. A Schematic diagram of experimental processes upon drugs addiction. B, C Infected WRD cells were treated with VER155008 (B) and quercetin (C) under indicated concentrations at 4 different time points according to drugs addiction, respectively. The viral protein levels were assessed with Western blot analyses. D-F MVC-infected WRD cells were treated with drugs at the IV time point of drug-addiction. MVC mRNA (D), vDNA (E) and secreted virions levels (F) were monitored and evaluated by qRT-PCR analyses. G, H Inhibitors were added at 24 h after MVC infection. vDNA (D) and virus particles (E) were detected via qRT-PCR. Data information: In (B-H), data shown in each panel are representative of three independent experiments. Statistical significance was determined with paired t-test (B, C) and two-way ANOVA (D-H). Asterisks indicate significant differences (\**P* < 0.05, \*\**P* < 0.01, \*\*\**P* < 0.001). Error bars represent standard error of the means (SEM).

Next, we added the drugs at 24 hpi and monitored MVC replication and extracellular virus production for an additional 24 h. At this point, viral replication compartments in infected cells are already established. Nevertheless, the addition of the compounds at 24 hpi still had strong and immediate inhibitory effect on vDNA (Fig. 8G) and extracellular secreted virions (Fig. 8H) in these conditions. Taken together, the results indicated that Hsp70s are required to facilitate MVC replication, both in its initial stage and its post-establishment. Not only, these inhibitors strikingly rendered virus progeny secretion failure in a time-dependent manner. The fact that the two chemically distinct compounds block MVC propagation is strong evidence that Hsp70s play vital roles in MVC infection. Despite their divergent roles in virus infection, Hsc70 and Hsp70 are surely involved in nearly all phases of the viral life cycle.

## DISCUSSION

In the course of viral life cycle, large amounts of viral proteins are synthesized in a short period of time, which can become a big challenge for viruses. Therefore, Hsp70s are hijacked by viruses to circumvent this limitation (Pujhari et al., 2019, Rosenzweig et al., 2019). Recently, accumulating evidence supports Hsp70s are a group of vital host cellular proteins that participate in proliferation of different kinds of viruses (Khachatoorian, Ganapathy et al., 2014, Khachatoorian, Ruchala et al., 2015, Taguwa et al., 2019). In this study, we investigated and dissected the roles of Hsc70 and Hsp70 in MVC propagation. Our results revealed that the Hsp70s chaperones are nearly involved in all phases of MVC life cycle, including viral entry, transcription, translation, replication, and virus production, which are correlated with their functions in folding and transporting newly synthesized proteins and promoting the disassembly of misfolded proteins.

For the first time, using MS analyses, Hsp70s were identified as targets interacting with NS1. We also demonstrated in vivo that both of Hsc70 and Hsp70 interact with NS1 and VP2, which happened completely independent of nucleic acid since the addition of DNase I did not abolish their interactions compared to NP1. Interestingly, we further showed both the nucleotide binding domain (N) and the substrate binding domain (C) of the two Hsp70s interact with NS1 and VP2, suggesting that both functional domains are required for the efficient collaborations between NS1/VP2 and Hsc70/Hsp70 during MVC infection. Furthermore, observation via immunofluorescence-based confocal microscopy revealed that localizations of Hsc70/Hsp70 and NS1 were more likely in the nucleus and localization with VP2 were mostly in the cytoplasm. These findings are in agreement with their roles in MVC, where NS1 is replication-associated and VP2 is involved in forming virus capsids. Infection of animal cells by viruses often results in elevation and re-localization of heat shock proteins, depending on the type of virus and host cell (Duncan, Cheetham et al., 2015). Increasing levels of hsp70 parallels the concentration of the insoluble viral coat protein in tobacco mosaic virus and other viral proteins in turnip mosaic virus and turnip crinkle virus, indicating the potent induction ability of viral proteins for chaperones including hsp70 (Jockusch, Wiegand et al., 2001). Here, a stimulation in expression of Hsp70 upon MVC infection was observed both in immunofluorescence assay and Western blot analysis. Additionally, transfection of viral protein-expressing plasmids also contributes to markable upregulations of Hsp70. We reasoned that these increasements were possibly specific to MVC infection since another member of Hsp family, Hsp90, remained nearly unchanged in infected cells.

We determined the activity of Hsc70 knockdown in the MVC cell culture system utilizing the shRNA-mediated gene silencing technique. Knockdown of Hsc70 resulted in a significant decrease in viral transcription, translation, replication and virus production. It should be noted that, interestingly, a higher stimulation in Hsp70 levels was observed under the condition of relatively less Hsc70. Considering the significant homology between Hsc70 and Hsp70, we speculated that the upregulation of Hsp70 might attribute to its potential compensatory role with Hsc70 in order to maintain cellular integrity during metabolic challenges. As such, the shHsc70-1 cells with higher levels of induced Hsp70 caused mild reduction while the shHsc70-2 cells with slight amounts of induced Hsp70 led to strikingly decrease in many aspects of MVC life cycle. In this situation, seemingly, Hsp70 complements with Hsc70 and plays a positive role during infection. Hence, we asked whether silencing of Hsp70 displayed same effects as Hsc70. An astonishing finding was that, distinct to what we have observed in Hsc70 knockdown scheme, specific reduction of Hsp70 resulted in evident growth in viral protein levels, transcription, replication, and virus production. Not only that, but heat shock (HS) treatment also significantly suppressed viral translation and virus production at 48 hpi, suggesting that Hsp70 might play a negative role upon MVC infection. However, forced overexpression of Hsp70 following different amounts of pFlag-Hsp70 transfection unveiled us for the first time that Hsp70 could exert dual effects in MVC life cycle. When Hsp70 was slightly overexpressed by 2μg, modest growth was seen in viral protein levels, whereas when Hsp70 was sufficiently overexpressed by 6μg, viral translation was visibly inhibited. This means that MVC infection is extremely sensitive to the content of cellular Hsp70. In other words, the effect of Hsp70 on viral infections is not inflexible. Within a small range of rise, Hsp70 acts as a promotive factor in viral life cycle, nevertheless, it exerts a strong inhibitive effect on MVC propagation when its amount is beyond the threshold, indicating that there must be a comparatively narrow window in which Hsp70 levels is confined.

It is not rare that Hsc70 and Hsp70 play distinct roles in virus infection. Interestingly, in the Hsp70 family, compared with the steadily expressed Hsc70, the role of inducible Hsp70 in viral infection is more complicated. Sometimes Hsp70 bears synergistic effects with Hsc70 as a positive regulator in viral life cycle. During hepatitis C virus (HCV) infection, non-structural protein NS5A interacts with both of Hsc70 and Hsp70. Individual knockdowns of Hsc70 and Hsp70 results in significant reduction of both intra- and extracellular virus production (Khachatoorian et al., 2014). In particular, Hsc70 is involved in the formation of infectious virions and plays roles in the budding and entry processes of the HCV lifecycle, but has no effects on intracellular HCV RNA replication (Khachatoorian et al., 2015). While Hsp70 positively regulates HCV replication by increasing levels of the replicase complex via enhancing HCV internal ribosome entry site (IRES) activity, possibly through its ATPase domain (Chen, Chen et al., 2010). However, sometimes the inducible Hsp70 acts as a negative modulator in virus infection. In the case of rotavirus, knockdown of Hsp70 by siRNA resulted in increased viral structural proteins and rotavirus morphogenesis in Caco-2 cell line through a ubiquitin-dependent process (Broquet, Lenoir et al., 2007). The Hsp70 family in yeast comprised of nine members, encoded by four SSA genes, two SSB genes, two SSE genes, and SSZ1. It has been reported that deletions of both SSA1 and SSA2 resulted in a significant reduction in flock house virus (FHV) RNA accumulation, however, deletion of SSZ1 or deletions of both SSB1 and SSB2 markedly enhanced FHV RNA and FHV protein A accumulation (Weeks, Shield et al., 2010). This suggests that, though with a high similarity in sequences, functions of Hsc70 and Hsp70 might differ from each other.

Based on their post-infection distinctions, we wonder whether they participate and differ with each other as well in the entry stage of MVC life cycle. Studies about roles of Hsc70 and Hsp70 in virus entry have been reported these years. Despite Hsp70s function mainly as cytoplasmic chaperones, they also act as virus receptors during many virus infections, including JEV, ZIKA and rotavirus (Broquet et al., 2007, Nain, Mukherjee et al., 2017, Pérez-Vargas, Romero et al., 2006, Pujhari et al., 2019, Taguwa et al., 2019). Interestingly, they perform differently in different permissive cell lines. In Huh7 cells, Hsp70, but not Hsc70 or Grp78, is crucial for the JEV entry possibly through interacting with viral E proteins, revealed by antibody blocking and siRNA knockdown experiments (Nain, Abdin et al., 2016). While in Neuro-2a cells, however, Grp78 interacts with JEV E-DIII and plays multiple roles in both entry and post-entry steps of JEV infection (Nain et al., 2017). In our hands, Hsc70 and Hsp70 both were detected on WRD cell membrane. Knockdown of Hsc70 and Hsp70 resulted in decrease and increase of intracellular viral loads as tested by viral proteins at 6 hpi and vDNA at 2 hpi, respectively. HS methods also lead to similar reduction on both aspects. Note that both of Hsc70 and Hsp70 interact with MVC VP2, we reasoned that Hsc70 and Hsp70 on cell membrane might interfere with the entry phase of MVC infection. As such, we wondered whether the results we obtained (Fig. 7, 8, 9) were interfered by the impacts of Hsc70 and Hsp70 on viral entry. Experiments of Hsp70 overexpression post infection and pI-MVC transfection showed that viral proteins and virions secretion were still modulated, indicating Hsc70 and Hsp70 are nearly required for all phases in MVC life cycle.

While we have observed the distinctions of viral protein levels upon different Hsp70 transfections (2μg, 6μg), the mechanisms under the regulation of Hsp70 on viral proteins remained unclear. It is well established that Hsp70 is a crucial regulator of CHIP-mediated ubiquitination and degradation. CHIP is a chaperone-dependent E3 ligase able to ubiquitinate misfolded proteins for proteasomal degradation with the help of the Hsp70 chaperone machinery. Hence, Hsp70-CHIP axis is a vital mechanism for maintaining proteostasis. This correlates well with our findings that during MVC infection, NS1 and VP2 are ubiquitinated, which is strongly related to the levels of Hsp70 and their degradation. We also show here that the addition of proteasome inhibitor MG132 induced elevations in the NS1 and VP2 levels. These are firm evidence that Hsp70 regulates viral proteins through a ubiquitin-dependent degradation pathway. However, whether the manipulation of viral proteins is involved only in this Hsp70-dependent ubiquitination pathway remains to be clarified. We speculated that specific depletion of inducible Hsp70 could possibly lead to compensatory upregulations of other isoforms which in turn might be recruited by MVC. These two views can coexist without contradiction.

Taking the intriguing circumstance into consideration, we took advantage of two known inhibitors in the following experiments, quercetin and VER155008, both of which have been reported widely in many virus infections. Quercetin works mainly on the synthesis of Hsp70s, with preferentially inhibitory effect on the inducible Hsp70 than the constitutive Hsc70 (Ogawa et al., 2014). In rabies, treatment with quercetin resulted in a significant decrease of viral mRNAs, proteins, and particles (Lahaye, Vidy et al., 2012). The folding action of Hsp70s is coupled to the ATPase activity. Adenosine derivative compound VER155008, as an ATP analog, induces conformational changes of Hsp70 family through competitive insertion into the ATP-binding pocket. The change in Hsp70s protein levels in response to VER155008 treatment may be cell type or context-dependent. This structure-based ATP-competitive modulator markedly inhibited the synthesis of baculovirus proteins, genome replication and the production of budded virions (Shrestha, Patel et al., 2016). Both inhibitors are implemented at non-toxic concentration windows for WRD cells. An advantage of these compounds over knockdown approach is that they can block the action of multiple Hsp70 isoforms and likely leave other chaperone functions intact. Here, time-of-addiction assay suggests the best effect of drug inhibition upon both Hsp70s and viral protein quantities is at a post-infection step. Results showed quercetin nearly abrogated virus transcription, replication and virion production synthesis compared to control group at multi-timepoint. VER155008 also suppressed MVC infection, but to a less extent. This indicates that Hsp70s are responsible for the establishment of an effective viral replication state. When drugs were added at 24 hpi, a timepoint when replication machinery is assembled, the kinetics results of MVC infection showed that inhibitors still remarkably and immediately reduced the vDNA and production of MVC particles, suggesting an important role of Hsp70s at least in virions egressing phase. Notably, quercetin exerts better performance than VER155008 at each aspect. When the majority Hsp70 family was shut down simultaneously, MVC nearly lost its capacity in conducting a successful infection.

In summary, this work has, for the first time, preliminarily shed lights on a primary but essential mechanism between the cellular 70 kDa family and MVC, a typical member of bocaparvovirus family. We demonstrated that Hsc70, Hsp70 chaperone interact with MVC NS1, VP2 proteins and their divergent effects on MVC infection. Our results favor the idea that Hsc70 and Hsp70 may serve as host factors that are able to control the amounts of viral proteins. These results are important for understanding the critical roles of Hsc70 and Hsp70 in the MVC viral life cycle, they also illustrate the significance of maintaining the proper balance of constitutive and stress-induced chaperones. Regardless, inhibitors of Hsp70 family lead to potent attenuation at many aspects of infection life cycle, which represents a potential target for the development of antiviral strategies.

## Materials and Methods

### Virus and cell culture

The Walter Reed canine cell/3873D (WRD) cell line and MVC were generously gifted from Professor Jianming Qiu of the Medicine Department of Microbiology, University of Kansas. Cos-1 and HEK293T (293T) cells were stored in our lab. Cells were cultured in Dulbecco’s modified Eagle’s medium (DMEM, Gibco) supplemented with 10% fetal bovine serum (FBS, TransGen Biotech) and 1% penicillin/streptomycin in 5% CO_2_ at 37 °C. For infection, WRD cells were seeded in culture dishes at a density of 20,000 cells/cm2. MVC was routinely propagated in WRD cells at MOI=10 (unless noted).

### Plasmids

Hsc70/Hsp70 were subcloned into pXJ40-Flag (pFlag) and pEGFP-C1 (pEGFP) vectors, VP2 was subsequently subcloned into pFlag and pCDNA vectors. NP1 was subcloned into pXJ40-HA (pHA) vector. pHA-NS1 was gift ed from Dr. Guan’s lab of the Wuhan Institute of Virology (WIV), Chinese Academy of Science (CAS). Primers used are as follows: pFlag-Hsc70 (F: 5’-CC CAAGCTTATGTCTAAGGGACCTGCAG-3’; R: 5’-TCCCCCGGGCTATCAACCTCTTCAATGGTAGGCCCAGAG-3’); pFlag-Hsp70 (F: 5’-CGCGGATCCATGGC TAAGAGCGCGGCCATCGGCATC-3’; R: 5’-TCCCCCGGGCTAATCCACCTCCTCGATGGTGGGGC-3’); pFlag-VP2 (F: 5’-CGCGGATCCGGTCGGAGCTTCTGGAGCTGAC-3’; R: 5’-TCCCCCGGGTTACAGGACCCTGTTTATTCCCC-3’); pCDNA-VP2 (F: 5’-CCGGAATTCATGGAACCGCAGGAAACTGAGAAC; R: CG CGGATCCTTACAGCGTTTTGTTTATGCCTC); pHA-NP1 (F: 5’-CGCGGATCC ATGTCTACGAGACATATGAGC-3’; R: 5’-TCCCCCGGGCTATTCGGAGGAGCCATCTAC-3’); pEGFP-Hsc70 (F: 5’-CCCAAGCTTCGATGTCTAAGGGACCTG CAG-3’; R: 5’-TCCCCCGGGTAGTACAAATTTGGGTCCTTCAAATGTG-3’); pEGFP-Hsp70 (F: 5’-CCGGAATTCTATGGCTAAGAGCGCGG-3’; R: 5’-TCCCCC GGGCTAATCCACCTCCTCGATGGTGGGC-3’); pEGFP-Hsc70 (N) (F: 5’-CCC AAGCTTTCCAGCACGGGAAAGTGG-3’; R: 5’-TCCCCCGGGTAAACCAGAAAGAATGGCTGC-3’); pEGFP-Hsc70 (C) (F: 5’-CCCAAGCTTCCCAGGCAGCC ATTCTTTCTGG-3’; R: 5’-TCCCCCGGGAATCAACCTCTTCAATGGTAGGCC CAGAG-3’); pEFGP-Hsp70 (N) (F: 5’-CCGGAATTCTATGGCTAAGAGCGCGG-3’; R: 5’-TCCCCCGGGTAGCAGCAGCAGGTCC-3’); pEGFP-Hsp70 (C) (F: 5’-CCGGAATTCCGTGCAGGACCTGCTGCTG-3’; R: 5’-TCCCCCGGGCTAAT CCACCTCCTCGATGGTGGGC-3’). The empty vectors containing these tags w ere referred to as controls.

### Identification of the NS1/VP2-interacting proteins

WRD cells seeded on a 10 mm dish were infected with MVC for 48 h and lysed in NP-40 lysis buffer (Beyotime, P0013F) containing protease inhibitors. After incubating on ice for 15 min, cell lysates were clarified by centrifugation at 14,000 g for 15 min. Supernatants were transferred into new tubes of which the 10% were saved for input. The remained liquids were added with anti-NS1 or -VP2 or -IgG antibodies and incubated at 4°C for 4 h followed by addition of protein A/G PLUS-Agarose (Santa Cruz Biotech, sc-2003) and incubation overnight. Proteins bound to the beads were washed five times with pre-cooled PBST buffer and eluted by boiling at 95°C for 5 min. In preparation for high performance liquid chromatography-tandem mass spectrometry (HPLC-MS/MS), protein samples were resolved in 10% sodium dodecyl sulfate polyacrylamide gel electrophoresis (SDS-PAGE), stained with Coomassie brilliant blue for 2 h and then distained. The gels were sent for MS analysis at the Beijing Qinglian Baiao Biotechnology Co., LTD.

### Coimmunoprecipitation (Co-IP)

Transfected cos-1 cells or infected WRD c ells were collected and lysed in NP-40 lysis buffer (Beyotime, P0013F) contain ing protease inhibitors. Each sample contained at least 500 μg of a total protei n. 2000U/mL DNase I (New England Biolabs, M0303S) and 1M MgCl_2_ were added to cell lysis at 1:100 (v/v) and placed in a 37°C for 30-60 min. Optim um concentration of antibodies against NS1/VP2/NP1/Hsc70/Hsp70/HA/Flag/GFP /ubiquitin or normal IgG (as a negative control) was added to each sample and then incubated at 4°C for 4 hours followed by addition of protein A/G PLUS -Agarose (Santa Cruz Biotech, sc-2003) according to manufacturer’s instructions. After incubation overnight, proteins bound to the beads were washed five tim es with pre-cooled PBST buffer and eluted by boiling. Samples were then appl ied to a standard Western blot procedure.

### Western blotting

Sample products were applied to SDS-PAGE and the transferred to polyvinylidene fluoride (PVDF) membrane. Protein bands were blocked with 5% skimmed milk, detected by incubating with corresponding primary antibodies and second antibodies and visualized with Super ECL Prime (Seven Biotech). The following primary antibodies were used for Western blotting: anti-Hsc70 (ab19136; Abcam; WB: 1:1000), anti-Hsp70 (ab47455; Abcam; WB: 1:1000), anti-ubiquitin (10201-2-AP; Proteintech; WB: 1:500); anti-NS1 (WB: 1:1000), anti-NP1 (WB: 1:1000) and anti-VP2 (WB: 1:1500) were prepared in house in cooperation with Abmart Company; anti-HA (66006-1-Ig; Proteintech; WB: 1:1000; IP: 3 μg), anti-Flag (66008-3-Ig; Proteintech; WB: 1:1000; IP: 3 μg), anti-GAPDH (TA-08; ZSGB-BIO; WB:1:1000), anti-E-Cadherin (14-3249-80; Invitrogen; WB: 1:1000), goat-anti-mouse (ZB-2305; ZSGB-BIO; WB: 1:5000), goat-anti-rabbit (ZB-2301; ZSGB-BIO; WB: 1:5000), mouse-anti-rabbit (A25022; Abbkine; WB: 1:2000), goat-anti-mouse (A25012; Abbkine; WB: 1:2000) and mouse-anti-rabbit (A25122; Abbkine; WB: 1:2000).

### Confocal microscopy and Immunofluorescence assay (IFA)

WRD cells cultured on confocal dishes (glass bottom cell culture dish, 801001, NEST) were subjected to MVC infection. Cos-1 cells were cultured on confocal dishes and followed by co-transfection of pHA-NS1 or pCDNA-VP2 and pEGFP-Hsc70 or pEGFP-Hsp70 plasmids. 48 hours later, cells were fixed in pre-cooled 4% paraformaldehyde-PBS or 100% ice cold Methanol for 20 min, permeabilized for 3 min with 0.1% Triton X-100, and blocked for one hour with 3% BSA at room temperature. The following primary antibodies were used for IFA: anti-Hsc70 (ab19136; Abcam; IFA: 1:100), anti-Hsp70 (ab47455; Abcam; IFA: 1:100), anti-VP2 (IFA: 1:100), anti-NS1 (IFA: 1:100). After incubation at room temperature for 2h or 4 °C overnight, cells were washed with PBS and double stained with Alexa Fluor 647-(bs-0295G-AF647; Bioss; IFA: 1:100) and/or Alexa Fluor 488-conjugated (bs-0296G-AF488; Bioss; IFA: 1:100) secondary antibodies. Nuclei were counter-stained with 4-6-diamidino-2-phenylindole (DAPI, ZLI-9557; ZSGB-BIO) for 5 min. Localizations were imaged using an Olympus scanning confocal microscope and analyzed with FV10-ASW 4.2 Viewer software according to software instructions.

### Quantitative Real-Time PCR (qRT-PCR)

Total RNA was extracted using Trizol agent (Invitrogen, 15596026) and cDNA was generated from 1 μg of R NA using a reverse transcription kit (TransGen Biotech). The mRNA expression levels of respective genes were shown as a ratio compared with GAPDH in t he same sample by calculation of cycle threshold (Ct) value in qRT-PCR ampli fication, of which 1.5 μL of the diluted cDNA were used as template. Viral D NA was extracted using a Quick-DNA^™^ Viral Kit (ZYMO Research). The standard plasmid pI-MVC (Sun et al., 2009) gifted from Professor Jianming Qiu of the Medicine Department of Microbiology, University of Kansas, was quanti fied by spectrophotometric analysis (NanoDrop ND-1000) and converted into co py number. Ten-fold serial diluted plasmids were used in absolute quantification. The primer sequences are: Hsc70 (F: 5’-TCCTGCTCCTCGTGGTGTT-3’; R: 5’-AAGCGGCCCTTGTCGTTAG-3’); Hsp70 (F: 5’-GCACGGGCAAGGCCAACA A-3’; R: 5’-CGTCCTCCACCGCACTCTTCAT-3’); NS1 (F: 5’-GAAGAAGACAT AACAGGTGA-3’; R: 5’-AACAGTGGAGGACGATTG-3’). qRT-PCR was perfor med using a qPCRsoft (Analytik Jena AG, Germany) with a SYBR®Green PC R Master Mix Kit (Takara Bio), according to the manufacturer’s instructions. Viral DNA levels and extracellular virus copies were calculated using the standard curves of PCR amplification obtained from the serial dilutions of pI-MVC plasmid.

### Short hairpin (sh) RNA-mediated gene silencing

shRNAs specific to Hsc70 and Hsp70 gene used in the study are as follows: Hsc70 (shHsc70-1: 5’-GGAACGTGTTGATCTTTGATT-3’, shHsc70-2: 5’-GCTGATTGGACGTAGATTTGA-3’), Hsp70 (shHsp70-4: 5’-GCGACAGGGTGTCTGCCAAGA-3’, 5’-GCTCATCGGCCGCAAGTTTGG-3’). shHsc70-1 and shHsc70-2 cloned into the pLKO.1-TRC vector were gifted from Huanzhou Xu of Dr. Guan’s lab of WIV, CAS. shHsp70-4 and shHsp70-5 were respectively cloned into pGMLV-SC5 vector (Genomeditech, GM-Mc-01306). Stable knockdown cell lines were generated by lentiviral infection followed with puromycin selection. 293T cells were seeded and cultured in 60 mm dishes. When the density reached 60% to 70%, the packaging plasmids PSPAX, pMD2.G and the knockdown plasmids of interest were co-transfected into 293T cells, respectively. 72 h later, the harvested supernatants were filtered and collected and stored at −80 °C. WRD cells were infected with lentivirus-containing supernatant for 12h and in the presence of 1 μg/ml polybrene (H8761, Solarbio). 48 hours after infection, cells were selected under puromycin at 1 μg/ml.

### Isolation of low molecular weight (Hirt) DNA and Southern blot

Cells were cultured in 60 mm dishes and then infected with MVC at 37 °C for 2 h. Cells were washed with PBS and grown in fresh medium for 48 h. The cells were washed twice with PBS and lysed in 2% sodium dodecyl sulfate (SDS) followed by proteinase K (0.5 mg/mL) treatment. Hirt DNA was harvested and applied to 1% agarose gel and transferred to Hybond N^+^ membrane (Southern blot was carried out in Dr Guan’s lab (WIV, CAS)). The blots were hybridized with the MVC VP2 genome probe using the DIG High Prime DNA Labeling and Detection Starter Kit II (Roche) according to the manufacturer’s protocol. Signals were detected in the ChemiDoc™ MP imaging system (BIO-RAD).

### Focus forming assay (FFA)

The cells were inoculated into a transparent 96-well culture plate. After overnight culture at 37 °C (5% CO_2_), 100ul of continuous ten-fold diluted complete medium containing virus was added to each well (at least two to three repeated wells were needed for each sample). Cells were then subsequently subjected to IFA followed by counting the last two wells that fluorescence was observable at 48 hpi. The virus titer was calculated from the average fluorescence number. For example, if the numbers of focus forming unit (FFU) counted in the 1:100 diluted wells were 80 and 74, the numbers counted in 1: 1,000 diluted wells were 6 and 8, the virus titer was calculated as follows: [(80+74) *100+(6+8) *1,000]/4/100 * 1,000 = 7.35* 10^4^ FFU/ml.

### Heat shock (HS) and inhibitors treatment conditions

For heat shock: WRD cells were grown to 60–70% confluence and then treated with heat shock (30 min at 42 °C) to induce heat stress responses. Cells were recovered at 37 °C for 9 hours before MVC infection; for experiments post infection, cells were infected and treated with HS at 1 hpi. For inhibitors: Quercetin and VER155008 (APE x BIO Technology, both) inhibitors dissolved in dimethyl sulfoxide (DMSO; Sigma, D8372) were performed under non-toxic concentrations respectively at 40 μM and 2 μM for the indicated times. For inhibitor: MG132 dissolved in DMSO was performed under concentration at 0.1 μM for the last 3 hours of MVC infection.

### Cell viability

WRD cells were cultured in detachable 96-well culture plates (TCP011896; BIOFIL) and grown in DMEM with 10% FBS at 37 °C for 12 h. After different length of time (30, 10, 5 and 1 min, respectively) upon HS treatment at 42 °C, cells were recovered to 37 °C for up to 48 h. The impact of HS on cells was evaluated by cell counting kit-8 (CCK-8; TransGen Biotech) assay, according to the manufacturer’s instruction. 10 μl of CCK-8 reagent was added to 100 μl DMEM each well. After 1-3 hours of incubation at 37 °C, the absorbance was measured at 450 nm.

### Virus entry

To determine the existence of Hsc70 and Hsp70 on cell membrane, a Membrane Protein Extraction kit (310005, BestBio) was recruited. Extracted proteins were subsequently applied to Western blot analysis. Endocytosed viral load in cells was detected at 2 hpi for viral DNA (vDNA) and 6 hpi for viral proteins. Cells were incubated with MVC at an MOI=20 on ice for 1 h, followed by one PBS wash and a shift to 37 °C for 2 h or 6 h with complete medium. After incubation, the cells were washed with chilled PBS and vDNA and proteins were then extracted. An addition of proteinase K treatment (1 mg/ml) for 20 min at 4 °C before vDNA extraction would allow any surface-bound virions that had not been internalized to be removed. The efficacy of the proteinase K treatment was established.

### Statistical analysis

Statistical significance was determined with t-test, one-way ANOVA and two-way ANOVA in Prism5 (GraphPad). Asterisks in figures indicate significant differences (\**P* < 0.05, \*\**P* < 0.01, \*\*\**P* < 0.001). Error bars represent the standard error of the means (SEM) for three independent experiments unless indicated.

## Acknowledgements

We are thankful to Professor Jianming Qiu (Medicine Department of Microbiology, University of Kansas) for providing WRD cells, pI-MVC plasmids, bocavirus MVC and constructive opinions. We are indebted to Wuxiang Guan (Wuhan Institute of Virology, Chinese Academy of Science) for the technical assistance. We thank members of the lab for advice on the figures. We thank Huanzhou Xu for the generous contribution of shHsc70 plasmids. All authors read and approved the final manuscript. This work was supported by the National Natural Science Foundation of China (NSFC, No. 31760041), the Key Research & Development Program of Ningxia Province (No. 2022BEG03112) and the Natural Science Foundation of Ningxia Province (No. 2021AAC02010).

## References

Binn LN, Lazar EC, Eddy GA, Kajima M (1970) Recovery and characterization of a minute virus of canines. Infection and immunity 1: 503–8

Broquet AH, Lenoir C, Gardet A, Sapin C, Chwetzoff S, Jouniaux AM, Lopez S, Trugnan G, Bachelet M, Thomas G (2007) Hsp70 negatively controls rotavirus protein bioavailability in caco-2 cells infected by the rotavirus RF strain. Journal of virology 81: 1297–304

Chen YJ, Chen YH, Chow LP, Tsai YH, Chen PH, Huang CY, Chen WT, Hwang LH (2010) Heat shock protein 72 is associated with the hepatitis C virus replicase complex and enhances viral RNA replication. The Journal of biological chemistry 285: 28183–90

Chromy LR, Pipas JM, Garcea RL (2003) Chaperone-mediated in vitro assembly of Polyomavirus capsids. Proceedings of the National Academy of Sciences of the United States of America 100: 10477–82

Deng X, Yan Z, Luo Y, Xu J, Cheng F, Li Y, Engelhardt JF, Qiu J (2013) In vitro modeling of human bocavirus 1 infection of polarized primary human airway epithelia. Journal of virology 87: 4097–102

Duncan EJ, Cheetham ME, Chapple JP, van der Spuy J (2015) The role of HSP70 and its co-chaperones in protein misfolding, aggregation and disease. Sub-cellular biochemistry 78: 243–73

Fernández-Fernández MR, Valpuesta JM (2018) Hsp70 chaperone: a master player in protein homeostasis. F1000Research 7

Flatt JW, Butcher SJ (2019) Adenovirus flow in host cell networks. Open biology 9: 190012

Glotzer JB, Saltik M, Chiocca S, Michou AI, Moseley P, Cotten M (2000) Activation of heat-shock response by an adenovirus is essential for virus replication. Nature 407: 207–11

Hu C, Yang J, Qi Z, Wu H, Wang B, Zou F, Mei H, Liu J, Wang W, Liu Q (2022) Heat shock proteins: Biological functions, pathological roles, and therapeutic opportunities. MedComm 3: e161

Jockusch H, Wiegand C, Mersch B, Rajes D (2001) Mutants of tobacco mosaic virus with temperature-sensitive coat proteins induce heat shock response in tobacco leaves. Molecular plant-microbe interactions: MPMI 14: 914–7

Khachatoorian R, Ganapathy E, Ahmadieh Y, Wheatley N, Sundberg C, Jung CL, Arumugaswami V, Raychaudhuri S, Dasgupta A, French SW (2014) The NS5A-binding heat shock proteins HSC70 and HSP70 play distinct roles in the hepatitis C viral life cycle. Virology 454-455: 118–27

Khachatoorian R, Ruchala P, Waring A, Jung CL, Ganapathy E, Wheatley N, Sundberg C, Arumugaswami V, Dasgupta A, French SW (2015) Structural characterization of the HSP70 interaction domain of the hepatitis C viral protein NS5A. Virology 475: 46–55

Kityk R, Kopp J, Sinning I, Mayer MP (2012) Structure and dynamics of the ATP-bound open conformation of Hsp70 chaperones. Molecular cell 48: 863–74

Lahaye X, Vidy A, Fouquet B, Blondel D (2012) Hsp70 protein positively regulates rabies virus infection. Journal of virology 86: 4743–51

Li F, Zhang Q, Yao Q, Chen L, Li J, Qiu J, Sun Y (2014) The DNA replication, virogenesis and infection of canine minute virus in non-permissive and permissive cells. Virus research 179: 147–52

Liu WW, Yang P, Chen XM, Xu DL, Hu YH (2014) Cloning and expression analysis of four heat shock protein genes in Ericerus pela (Homoptera: Coccidae). Journal of insect science (Online) 14

Majumder K, Wang J, Boftsi M, Fuller MS, Rede JE, Joshi T, Pintel DJ (2018) Parvovirus minute virus of mice interacts with sites of cellular DNA damage to establish and amplify its lytic infection. eLife 7

Nain M, Abdin MZ, Kalia M, Vrati S (2016) Japanese encephalitis virus invasion of cell: allies and alleys. Reviews in medical virology 26: 129–41

Nain M, Mukherjee S, Karmakar SP, Paton AW, Paton JC, Abdin MZ, Basu A, Kalia M, Vrati S (2017) GRP78 Is an Important Host Factor for Japanese Encephalitis Virus Entry and Replication in Mammalian Cells. Journal of virology 91

Ogawa Y, Sugawara S, Tatsuta T, Hosono M, Nitta K, Fujii Y, Kobayashi H, Fujimura T, Taka H, Koide Y, Hasan I, Matsumoto R, Yasumitsu H, Kanaly RA, Kawsar SM, Ozeki Y (2014) Sialyl-glycoconjugates in cholesterol-rich microdomains of P388 cells are the triggers for apoptosis induced by Rana catesbeiana oocyte ribonuclease. Glycoconjugate journal 31: 171–84

Pérez-Vargas J, Romero P, López S, Arias CF (2006) The peptide-binding and ATPase domains of recombinant hsc70 are required to interact with rotavirus and reduce its infectivity. Journal of virology 80: 3322–31

Pollock RV, Coyne MJ (1993) Canine parvovirus. Vet Clin North Am Small Anim Pract 23: 555–68

Pujhari S, Brustolin M, Macias VM, Nissly RH, Nomura M, Kuchipudi SV, Rasgon JL (2019) Heat shock protein 70 (Hsp70) mediates Zika virus entry, replication, and egress from host cells. Emerging microbes & infections 8: 8–16

Reyes-Del Valle J, Chávez-Salinas S, Medina F, Del Angel RM (2005) Heat shock protein 90 and heat shock protein 70 are components of dengue virus receptor complex in human cells. Journal of virology 79: 4557–67

Rosenzweig R, Nillegoda NB, Mayer MP, Bukau B (2019) The Hsp70 chaperone network. Nature reviews Molecular cell biology 20: 665–680

Shao L, Shen W, Wang S, Qiu J (2021) Recent Advances in Molecular Biology of Human Bocavirus 1 and Its Applications. Frontiers in microbiology 12: 696604

Shrestha L, Patel HJ, Chiosis G (2016) Chemical Tools to Investigate Mechanisms Associated with HSP90 and HSP70 in Disease. Cell Chem Biol 23: 158–172

Sun Y, Chen AY, Cheng F, Guan W, Johnson FB, Qiu J (2009) Molecular characterization of infectious clones of the minute virus of canines reveals unique features of bocaviruses. Journal of virology 83: 3956–67

Taguwa S, Maringer K, Li X, Bernal-Rubio D, Rauch JN, Gestwicki JE, Andino R, Fernandez-Sesma A, Frydman J (2015) Defining Hsp70 Subnetworks in Dengue Virus Replication Reveals Key Vulnerability in Flavivirus Infection. Cell 163: 1108–1123

Taguwa S, Yeh MT, Rainbolt TK, Nayak A, Shao H, Gestwicki JE, Andino R, Frydman J (2019) Zika Virus Dependence on Host Hsp70 Provides a Protective Strategy against Infection and Disease. Cell reports 26: 906–920.e3

Wan Q, Song D, Li H, He ML (2020) Stress proteins: the biological functions in virus infection, present and challenges for target-based antiviral drug development. Signal transduction and targeted therapy 5: 125

Weeks SA, Shield WP, Sahi C, Craig EA, Rospert S, Miller DJ (2010) A targeted analysis of cellular chaperones reveals contrasting roles for heat shock protein 70 in flock house virus RNA replication. Journal of virology 84: 330–9

Wu HH, Wang B, Armstrong SR, Abuetabh Y, Leng S, Roa WHY, Atfi A, Marchese A, Wilson B, Sergi C, Flores ER, Eisenstat DD, Leng RP (2021) Hsp70 acts as a fine-switch that controls E3 ligase CHIP-mediated TAp63 and ΔNp63 ubiquitination and degradation. Nucleic acids research 49: 2740–2758

Zhang X, Yu W (2022) Heat shock proteins and viral infection. Frontiers in immunology 13: 947789

